# Acoustic monitoring for tropical insect conservation

**DOI:** 10.1101/2024.07.03.601657

**Authors:** Klaus Riede, Rohini Balakrishnan

## Abstract

Monitoring the species-specific sounds produced by insects could provide us with a rapid, reliable, non-invasive measure of tropical ecosystem health and biodiversity. Although acoustic biodiversity monitoring has made rapid progress over the past decade, the focus has been mostly on vertebrates, even though insects far outnumber them, and tropical soundscapes are dominated by insect sounds. Here we provide an overview of song features for the major sound-producing insect groups, identify technological milestones and describe impediments for analyzing tropical soundscapes and insect communities. We review some promising best-practices using singing insects for non-invasive acoustic profiling and tracking of diversity in rainforest ecosystems under threat. We suggest a roadmap for joint research efforts to accelerate acoustic assessments of singing insects based on re-using the wealth of existing data from Passive Acoustic Monitoring (PAM) in combination with curated multimedia repositories and citizen science.

## Introduction

Tropical forest soundscapes are dominated by insect voices (Aide 2017)! Due to the high number of insect species in tropical rainforests (Stork 2018), many of the ambient sounds are generated by these mostly invisible but audible creatures. Even though most insect groups are silent, sound-producing crickets, katydids and cicadas, together with some less audible groups such as moths and mosquitoes surpass vertebrates in species numbers and abundance. In addition, many insect species can only be found in the forest canopy or are active at night, posing a challenge to sampling by humans. Therefore, non-invasive acoustic monitoring is an ideal method to assess insect diversity in species-diverse tropical ecosystems. Even though tropical forests buffer warming due to climate change by providing a cooler microclimate, slight increases in temperature might affect tropical insects, which seem to be adapted to narrow climate niches (Chowdhoury 2023). The power of ecoacoustics to monitor climate change effects on biodiversity (Krause and Farina 2016) could be potentiated by using insects as proxies.

However, the focus of human attention is without doubt on vertebrates, both in conservation (Titley et al. 2017, Caldwell et al, 2024) and bioacoustics (see this volume). Here we present an overview of these hidden and overlooked contributors to present-day, but also prehistoric (Rust et al. 1999, Gu et al. 2012) soundscapes. These include sounds beyond the human hearing range, reaching far into the ultrasound (Sarria-S et al. 2014), but also into the vibrosphere. This acoustic diversity is a major challenge, and it may seem like a mission impossible to track individual songsters within complex tropical soundscapes (Fig. 1).

**Figure 1.**
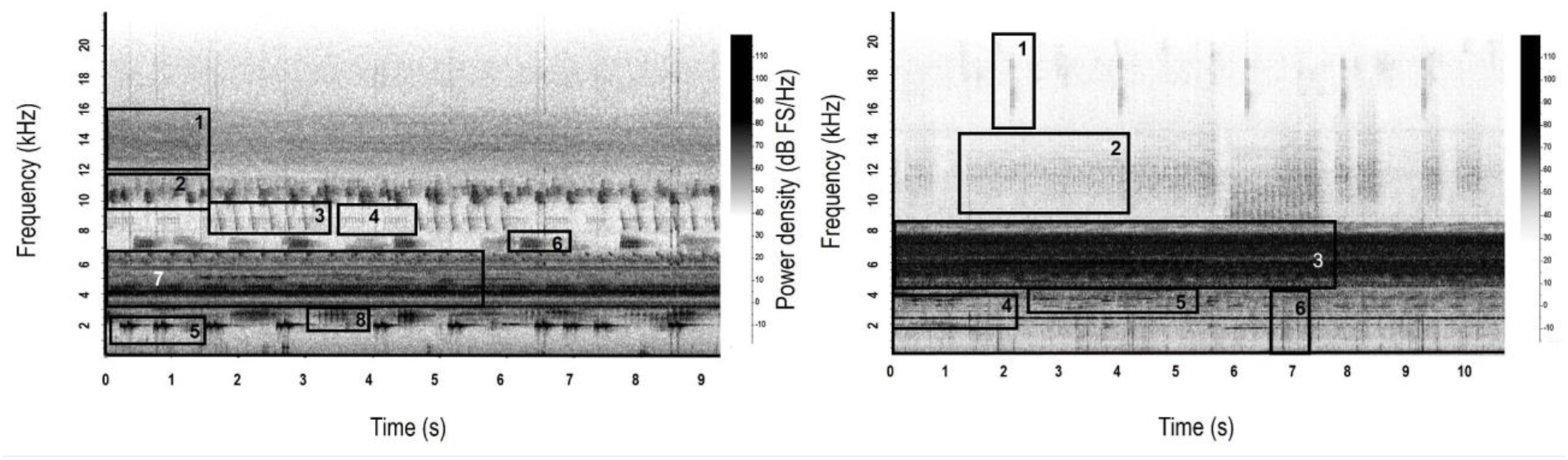
Distinguishing between gryllid, katydid, cicada and frog calls can be difficult in tropical rainforest soundscapes. Spectrograms of nocturnal rainforest soundscapes from India (Silent Valley, Kerala, May 2001: left panel) and Ecuador (San Pablo de Kantesya, Sucumbios,, January 2009: right panel). Numbered boxes show calls of species from different taxonomic groups. Left panel: Boxes 1-5: katydids, including a low-frequency tonal species (5); box 6: gryllid or katydid? (7-8 kHz); box 7: multiple gryllid species (3-7 kHz); box 8: tree frog. Right panel: Boxes 1-2: katydids; box 3: multiple gryllid species; box 4: low frequency tonal cicada (2.5 kHz); box 5: frog (3.5 kHz); box 6: water drops (rain). Spectrogram settings: Hann window, 1024 samples length, 90% overlap. Original wav files available in Suppl. Material.

In addition, inadequate knowledge of insect taxonomy delays biodiversity assessments, due to the high number of undescribed species (Stork 2018) and an acute taxonomic impediment (Löbl et al. 2023). We have considerable methodological problems in attributing identified songs (sonotypes, ethospecies) to respective species: most sonotypes lack voucher specimens, and most museum specimens lack sound recordings. Consequently, there is now a whole branch of ecoacoustics relying more on overall acoustic indices than on identification of well-defined songs and species.

Here we provide an overview of sound production and song features for the major sound-producing insect groups, identify technological milestones facilitating bioacoustic research in the tropics and review recent innovative approaches for analyzing insect communities contributing to tropical soundscapes.

While there are preliminary promising results to use such non-invasive assessments for insect conservation in anthropogenically threatened rainforest ecosystems, the method is still far from well established. Therefore, we suggest a roadmap for joint research efforts to accelerate acoustic assessments of singing insects, based on re-using the wealth of existing data from Passive Acoustic Monitoring (PAM, cf. Sugai et al. 2019, Gibb et al. 2019) in combination with curated multimedia repositories with references to voucher specimens, as well as pictures and videos provided by citizen scientists. In addition, considerable progress has been made in automatic song recognition, which allows identification of prominent songsters in large datasets from an increasing number of acoustic monitoring sites (Faiß and Stowell 2023; Madhusudhana et al. 2024).

### Section 2: Insect sounds: production, structure and diversity

Insects are classified into 29 orders, of which only a few produce sounds. Most prominent among them are the hundreds of species of crickets and grasshoppers (Orthoptera), belonging to an evolutionarily ancient lineage that evolved acoustic communication more than 250 million years ago (Song et al. 2020), and the cicadas (Order Hemiptera), that produce the familiar, loud choruses on hot summer days. Some species of arctiid and noctuid moths, and a few butterfly species (Order Lepidoptera) produce sounds as adults or larvae for communication or defense against predators (Low et al. 2021). Other groups known to produce sounds include several beetle groups (Order Coleoptera), mosquitos and flies (Diptera), bees and ants (Hymenoptera). The handbook by Drosopoulos and Claridge (2006) provides an excellent overview of insect sound production and its behavioural significance, including vibratory communication.

Sound production in different groups of insects typically happens using five basic mechanisms (Morris 1999; Low et al. 2021): flight tones or audible wing beats (mosquitoes, flies, bees), stridulation (rubbing one body part against another: crickets, katydids, grasshoppers, weta, ants), deformation of body parts (tymbals and crepitation: cicadas, some grasshoppers, arctiid and noctuid moths, butterflies), percussion (striking a body part on an external substrate) and tremulation (shaking body parts to send a vibrational signal through the substrate). Among these, vibrational signalling using tremulation or stridulation (as in ants) is probably ubiquitous among insects, but little explored in insect orders other than hemipterans (planthoppers) and neuropterans (lacewings). Characterizing and identifying species and behaviours based on these vibrational signals would enable sampling across diverse insect orders using vibroscapes in the future (Sturm et al. 2022). Given the limited number of insects whose vibrational signals have been characterised, however, and the technical challenges of recording vibroscapes, we limit our discussion to audio sampling and soundscapes.

The flight tones of mosquitos, flies and bees are of relatively low frequency (100-250 Hz for flies and bees, 300-700 Hz for mosquitoes) and species-specific, often carrying information on both the species and sex of the signaler (Kawakita and Ichikawa 2019, Kim et al. 2021, Rodríguez et al. 2024). Identifying and discriminating these acoustic signals should enable monitoring of populations and species of these insect groups which are globally important as disease vectors and/or providers of important ecosystem services such as pollination. The fact that most other insect groups produce sounds at higher frequencies should make the task easier (Fig. 1).

One of the most widespread modes of sound production is stridulation or rubbing different body parts against each other, generating sounds above 1 kHz (an exception is the Malaysian katydid *Tympanophyllum arcufolium* (Haan, 1843) with, at 600 Hz, the lowest known frequency call for a long-distance terrestrial insect signal: Heller 1995). Among the Orthoptera, ensiferan (crickets, katydids and wetas) signals in a soundscape span a large frequency range (600 Hz -150 kHz). They can be differentiated by their distinct frequency bands, lack of frequency modulation (as opposed to mammalian and bird calls), their stereotyped, repetitive nature and species-specific call temporal structures (Fig.1). Most true crickets (Grylloidea) produce tonal signals between 2-9 kHz, with diverse, species-specific temporal structures (Bennett-Clark 1998). In tropical forests, multiple species calling in this frequency band often co-occur and call together, making it challenging to discriminate their species-specific patterns, especially from single-microphone recordings (Fig. 1). In contrast, the calls of katydid (Tettigoniodea) species show an immense diversity of frequencies and bandwidths (Diwakar and Balakrishnan 2007, Tan et al. 2023), from low-frequency tonal calls similar to true crickets (the pure-tone Indo-Malayan false leaf katydid *Onomarchus uninotatus*: 2-3 kHz: Fig. 1), to calls of exceptional bandwidth (the Asian genus *Mecopoda* whose calls span a frequency range from 2-80 kHz or more: Nityananda and Balakrishnan 2006; Heller et al. 2021). The highest known frequency produced by an insect, a tonal call of 150 kHz, is by the katydid *Supersonus* spp. (Sarria S et al. 2014). Weta species of Southern India (Genus *Gryllacropsis*) produce loud, rasping, low frequency sounds (1.7 kHz: Diwakar and Balakrishnan 2006).

Most grasshoppers (Acridoidea: Acrididae) stridulate by rubbing their legs against the forewings: their calls are often broadband, at frequencies ranging from 2-40 kHz and often of less intensity than other orthopteran calls (Riede 1998). An interesting exception is the pneumorid or bladder grasshoppers of South Africa, the males of which produce extremely loud calls (by femoro-abdominal stridulation) that can be heard up to 2 km away (Römer et al. 2014). It should be noted however that most tropical forest grasshoppers do not produce sounds at all (Riede 1987) and therefore do not feature in tropical forest soundscapes. However, notable exceptions are some neotropical lubber grasshoppers (Acridoidea: Romaleidae), such as *Prionacris* species (*P. cantans* and *P*. cantrix: Descamps 1981).

Other insect groups (cicadas, noctuid and arctiid moths) produce sounds using tymbal organs, which rely on structural deformation followed by rapid energy release (Bennett-Clark 1998, Neil and Holderied 2021). The most conspicuous among these are the calls of cicadas, that show considerable overlap in structure and frequency space with katydids (Fig. 1: Ecuador). Cicada species’ songs can be characterized by temporal patterns and spectral features, including formants. Boulard (2006) created “Cards for Identification by Acoustics” and introduced the terms sonotypes and etho-sonotypes. Inspite of considerable sound intensities and typically diurnal or crepuscular song activity (Diwakar and Balakrishnan 2007), identification of songsters is difficult. Speech-like formants of cicada calls facilitate preliminary nomenclature with onomatopoetic names, which were used by Gogala and Riede (1995) to characterize the prominent songsters dominating the dusk chorus in Borneo (Malaysia). In contrast, most tropical crickets and katydids stridulate at night, and the timing of calling can be a simple yet effective way to separate these groups in soundscape recordings.

Some arctiid moths produce ultrasonic signals (Heller and Krahe 1994) very similar in structure to katydid calls and cannot easily be distinguished from katydids in soundscape recordings. Nakano et al. (2009) report on whispering sounds from moths, indicating that more mystery sounds can be expected in tropical soundscapes.

### Section 3: From bioacoustics to ecoacoustics

Insect bioacoustics has a long history and is fundamental for insect ecoacoustics. As early as 1854, Alexandre Yersin published musical scores of grasshopper songs, demonstrating their stereotyped, species-specific idiosyncrasies which are fundamental for species identification in soundscapes. During the 20th century focal recordings of individual species using tape recorders were stored in phonotheks which laid the ground for call libraries, including tropical species. In 1897, the physicist Amos Dolbear demonstrated the influence of temperature on cricket song pulse frequency, now known as Dolbear’s law. His publication about the “thermometer cricket” (Dolbear 1897) became widely popular and was used by the public in the US to measure temperature listening to a cricket and counting pulses with a stopwatch. This effect is highly relevant, particularly for song recognition algorithms, because reference recordings might vary considerably, and should be corrected for temperature effects.

The fascinating history of first voluminous tape recorders to the recent development of small, cheap digital recorders has been reviewed by Pavan et al. (2022). The invention of programmable autonomous recording units (ARUs) was the pre-requisite for the increasing number of present-day acoustic monitoring projects.

### Call libraries compiled from focal recordings

Determination of insect songsters requires a reference call library from identified species. Traditionally, such recordings were made with tape recorders either in the field or from captive individuals. Songs are published as sonograms, and ideally tapes are stored in phonotheks. In recent times, some of these historic recordings were digitized and made available in appropriate multimedia databases (Baker et al. 2015), but there is no centralized repository for insect sounds (cf. Riede 2018).

Morris et al. (1989) pioneered ultrasonic recording of katydids in Ecuador. Naskrecki (2000) published pictures and songs of katydids from Costa Rica, and ter Hofstede et al. (2020) published fact sheets including detailed sound descriptions for Panamanian katydids from Barro Colorado Island research station.

Braun (2002) made a thorough inventory of katydid songs from a montane rainforest in Southern Ecuador. Taxonomic description of new species from this site is still on-going (Braun 2021), and there are still several “mystery songs” which are probably from katydids, but lacking voucher specimens. Nischk (1999) compared cricket communities from Amazonian lowland rainforest site with a less diverse montane rainforest site in Ecuador (Nischk and Riede 2001). Some of the larger spider crickets (Phalangopsinae) and their songs were formally described (Nischk & Otte 2000), but only two of the smaller swordtail crickets (Trigonidiinae: Desutter-Grandcolas & Nischk 2000). About 30 sonotypes with voucher specimens are awaiting taxonomic description!

While there are some bioacoustic studies from the Neotropics which could serve as a starting point, comparable acoustic assessments are even rarer for the Paleotropics. The monumental work by Otte & Alexander (1983) on Australian crickets includes sonograms and song parameters for hundreds of newly described cricket species, including tropical Northern Australia. Some of these genera are widely distributed, occurring throughout the entire Australasian tropics, or are even pantropical. For some of these cricket species, song recordings were published on a CD included in Rentz (1996), together with songs of Australian katydids.

Hemp et al. (2023) reviewed the speciose East African katydid genus *Dioncomena*, describing several new species and their remarkable songs. Species descriptions including sound appear continuously but at a slow rate and often with remarkable delay: Heller & Helb (2024) described the song of the Brazilian *tanana* katydid 150 years after was noticed by Wallace and illustrated by Darwin.

Given that large numbers of insect species’ calls are unknown, especially in tropical forests of Asia and Africa, an alternative strategy has been to work with ‘sonotypes’ (unique call patterns) as proxies for species in soundscape recordings (Fig. 1). Since most insect species have stereotyped, species-specific calls, this may be a reasonable approach for species diversity estimation, especially in species-rich, poorly characterized and rapidly changing tropical landscapes typical of Asia and Africa. Such an approach was used by Grant (2014) to examine the acoustic insect assemblage of a rainforest in Borneo. To finally translate into effective monitoring and insect conservation, however, the sonotypes will have to be ground-truthed and assigned to species (as indicated in the flowchart in Fig. 2). Particularly low frequency insect sounds can easily be confounded with frogs or even birds – only recently, Dias et al. (2017) found that the katydid *Paracycloptera grandifolia* generates a hitherto enigmatic call from the canopy of Brazilian Atlantic rainforest.

**Figure 2.**
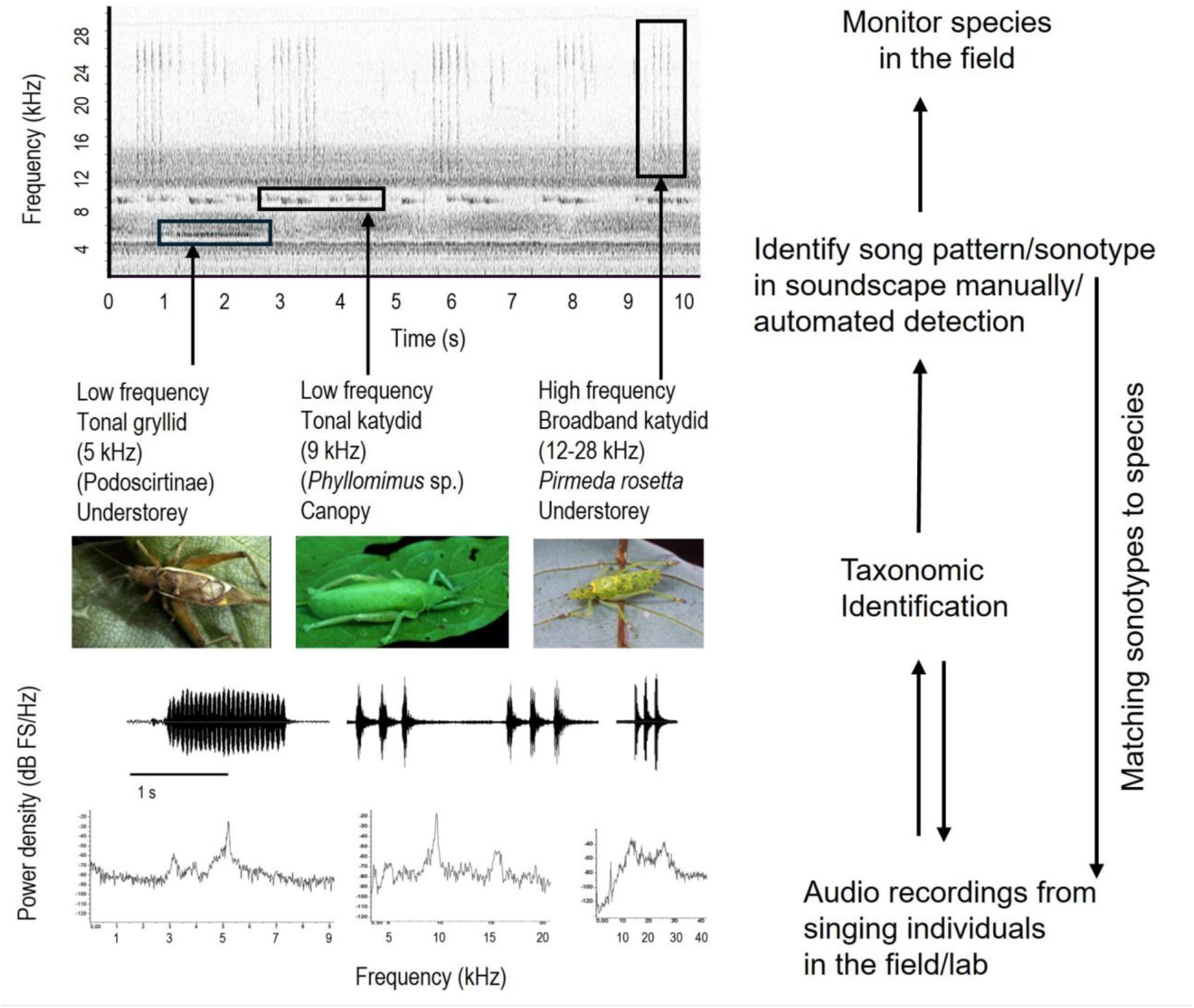
From song libraries to soundscapes and vice versa. Top left: Spectrogram of rainforest nocturnal soundscape from Kudremukh National Park, India (January 2005; settings: Hann window, 1024 samples, 90% overlap), with three species’ calls highlighted in boxes. Bottom left panel: Oscillograms and power spectra of the three highlighted species. Note that they have been classified up to different levels, illustrating the taxonomic impediment to species monitoring in tropical ecosystems. Right: Flow chart illustrating the iterative steps towards effective passive acoustic monitoring of insect species in the field.

### Acoustic monitoring of insect diversity at the species level

Accurate identification of singing insects in the habitat requires human experts and a call library attributable to individual voucher specimens in museum repositories. Trained human listeners could work in the field, with audio recordings, or both. This is the traditional and perhaps the most accurate way yet to acoustically monitor insect diversity in the audio range, comparable to bird counts (cf. Darras et al. 2018). Diwakar et al. (2007) monitored a rainforest insect community in southern India using a combination of point counts by a trained human listener and simultaneous audio recordings. The calls and identities of the insect species were established earlier by making audio recordings of individual insect callers which were then captured for taxonomic identification (Fig. 2, Diwakar and Balakrishnan 2007). Both human and recorder had high probabilities (> 80%) of detecting broadband signals and tonal calls in the range of 3-8 kHz. The human listener could, however, also elucidate the number of individuals of each species, information that would be difficult to extract from single-microphone soundscape recordings.

In contrast, human listeners will be lost in many neotropical soundscapes, (Fig. 1: Ecuador) because most katydids there generate short ultrasound signals. In addition, the pronounced background “green noise” generated by crickets between 5 and 9 kHz masks other signals considerably. The only insect species reliably detectable for human listeners would be cicadas and crickets singing below 5 kHz, particularly spider crickets (Phalangopsinae, cf. Nischk & Otte 2000). However, direct assessment by human listeners in the Neotropics is possible in less complex soundscapes such as mountain forests (cf. Nischk and Riede 2001) or detecting prominent songsters such as the lubber grasshopper *Prionacris* spp during the day.

Symes et al. (2022) evaluated soundscape spectrograms generated by PAM to elucidate spatio-temporal patterns of calling activity in Barro Colorado rainforests, based on the call library by ter Hofstede et al. (2020), by determination of temporal and spectral parameters.

### Challenges for acoustic insect monitoring by PAM

Whilst the technical challenges of sampling across the broad frequency range of insect calls have recently been overcome with the development of compact yet powerful recorders such as AudioMoth (Hill et al. 2019), recognising and identifying the signals of different insect species remains a challenge. A major reason for this is the lack of reference call libraries for sound-producing insects, especially in tropical regions outside the Americas.

A second challenge for monitoring of insect, and indeed other taxon, diversity from soundscapes is the ‘big-data’ problem. Soundscape data are easy to collect, and this results in the accumulation of incredibly large quantities of data that would take decades to evaluate manually. This has led to a focus on developing methods for automated detection and recognition of species’ signals. Much of the focus of automated recognition has been on vertebrate taxa, especially birds and mammals. Recently however, automatic recognition algorithms have been developed to discriminate different mosquito and bee species from their flight sounds (Kawakita and Ichikawa 2019), and cicada, cricket and katydid species based on their calls (Noda et al. 2019, Tey et al. 2022, Faiß and Stowell 2023), some with impressive accuracies of species-level discrimination (90-98%). Most of the algorithms have however been developed using recordings of individual species available in databases, with data augmentation techniques to introduce noise. While these are a promising first step, the algorithms need to be tested on soundscape recordings in the field that are likely to be much noisier and, in some cases, have signals from multiple individuals and species in the same frequency bands (Fig. 1). In a recent study, Madhusudhana et al. (2024) used a variety of data augmentation techniques together with machine learning to explore automated recognition of the calls of 31 katydid species from field soundscape recordings on Barro Colorado Island, Panama. Given the diversity of insect species and soundscapes worldwide, such studies need to be replicated and the performance of automated algorithms evaluated and improved across the tropics. It should be noted that insect ears and nervous systems are well adapted to cope with complex soundscapes (cf. Römer 2020), which might inspire technical improvements in signal processing.

### From acoustic indices to unsupervised clustering

In a seminal paper Sueur et al. (2008) introduced acoustic indices based on soundscape entropy as a tool for rapid diversity assessment of vocalizing species. For recordings from Tanzanian dry lowland coastal forest patches, indices reflected lower species diversity in disturbed forests. However, the data do not reveal individual target species, as needed for conservation. Discussing the reliability of acoustic indices, Sethi et al (2023) conclude that “existing analytical approaches behave unpredictably” and Alcocer et al. (2022) go even further, stating: “After more than a decade of research, we still lack a statistically informed synthesis of the power of acoustic indices that elucidates whether they effectively function as proxies for biological diversity”.

Despite these limitations, PAM datasets generated during ecoacoustic assessments on the landscape level could be extremely useful for insect conservationists, if available for re-use. The recently published Worldwide Soundscape projects database https://ecosound-web.de/ecosound_web/collection/index/106 is an ideal tool to identify acoustic “biodiversity time capsules” (*sensu* Sugai and Llusia 2019) for re-analysis. Particularly promising are unsupervised learning techniques using clustering algorithms, which suggest clusters of sound types (Ulloa et al. 2018). Using clustering algorithms, Guerrero et al. (2023) were able to identify insect sonotypes in four datasets from Colombia, including two from extremely species-rich protected tropical forest areas. The study provides a detailed methodological section describing the algorithm and necessary pre-processing steps for feature extraction. An impressive number of approaches including ready-to-use software packages are now available for such advanced analysis, extensively reviewed by Napier et al. (2024). However, the IT (information technology) knowledge needed to use these sophisticated tools is still very high. A promising application for conservationists is provided by Arbimon: https://arbimon.org/, “an open source ecoacoustic analysis platform empowering scientists and conservationists with an efficient way to upload, store, and analyze mass amounts of acoustic data”. Arbimon offers several types of analyses via cloud computing, such as pattern matching based on only one template, supervised detection models and unsupervised clustering (https://arbimon.org/howitworks). Users can upload pattern templates for public use. Again, vertebrate patterns outnumber insects (as by 17/5/2024, 229 amphibian songs vs 9 insect patterns) but we predict rapid growth.

### Acoustic monitoring for conservation

Species discovery and monitoring are necessary but not sufficient for conservation. Examples for direct integration of ecoacoustics into tropical (insect) conservation projects are still scarce. Hugel (2012) used acoustic monitoring of selected cricket and katydid species as an indicator for regenerating forest vegetation on Rodrigues Island, where an endemic cricket community is associated with endemic island vegetation but threatened by invasive species. Removal of invasives led to regeneration of native forest, and acoustic recording was used to monitor successful regeneration of the orthopteran community.

Starting with identification of microendemisms in New Caledonia based on soundscapes, Gasc et al (2013) initiated a series of conservation monitoring activities based on acoustics. Anso et al. (2022) studied this cricket community as indicators of succession and found similar patterns when comparing silent species with those acoustically detectable. Warren et al. (2016) combined molecular with acoustical surveys to identify hotspots of cricket species radiation on tropical islands.

### Perspectives and a roadmap for future insect monitoring in the tropics

Despite some promising examples, tangible results for tropical insect conservation using acoustic monitoring are still scarce. Lack of sound reference libraries together with the taxonomic impediment and a lack of appropriate IT infrastructure (databases combining voucher specimens and song recordings) will be a challenge difficult to overcome in the near future.

Acoustic indices provide valuable general insights but are far too vague for insect monitoring on the species level, as needed for conservation. There is no shortcut avoiding dedicated sound inventories: Fig. 2 illustrates the iterative loop between soundscapes, sonotypes, individual recordings and species identification for effective monitoring and conservation of insects. The scarcity of such exemplary, “traditional” approaches providing reliable sound libraries is in stark contrast to the data tsunami generated by PAM recordings. Progress in automatic sound recognition is mind-boggling, but most biologists are overwhelmed by the IT literacy required for analysis of huge datasets (as reviewed in Kvsn et al. 2020).

An interesting approach to overcome the paucity of reference call libraries is to use predictive methods, wherein species’ call frequencies and temporal features such as pulse duration may be inferred using a combination of morphological and biomechanical measurements together with statistical and/or Finite Element Models (FEM). These methods have been validated in crickets and katydids (Montealegre-Z et al. 2017; Godthi et al. 2022) and shown to predict call frequencies with high accuracy. The method was also used to reconstruct the calls of rare and fossil orthopteran insects (*Prophalangopsis obscura*: Gu et al. 2012; *Archaboilus musicus*: Woodrow et al. 2022). This opens the possibility to predict call frequency and some temporal parameters from museum specimens. An extension of FEM modelling to other sound-producing insect groups would enable similar predictive modelling of calls from museum specimens. To be useful in ecoacoustic monitoring, however, approaches based on predictive modelling and/or sonotypes will need to be followed up with fieldwork in type localities, using soundscape recordings and ground-truthing to validate the presence of these species.

Global conservation policies require a strategy to obtain robust data for measuring progress against biodiversity targets and insect ecoacoustics needs to progress further to meet these objectives (Hochkirch et al. 2021). Once regional sound libraries have been established, acoustic assessments can be extended to wider geographic areas, eventually with help from citizen scientists. Strategic selection of certain sonotypes and close cooperation between citizen science and experts could help to accelerate generation of reliable data. For example, the prominent calls of neotropical *Prionacris* lubber grasshoppers are easy to record and identify, and probably are already present as “bycatch” in numerous Xeno-canto bird recordings from Ecuador and Peru, misidentified as bird calls, or classified as “mystery sounds” (https://xeno-canto.org/mysteries). Once resolved, mystery sound recordings could turn rapidly into validated data points appearing in GBIF (www.gbif.org). Inaturalist facilitates creation of dedicated projects (https://www.inaturalist.org/projects/new) allowing the pooling of data for a certain group of singing insects in an area, for example, Ecuadorian spider crickets (Phalangopsinae).

In many cases, identification of species generating these clearly structured, well-defined songs should be feasible, using the available sonograms and parameter datasets. In summary, citizen science has enormous potential, but close cooperation between scientists and citizen participants is needed to maintain scientific rigour (cf. Sheard et al. 2024).

Despite the challenges and knowledge gaps outlined here, and the difficulties of disentangling complex tropical multispecies soundscapes, the perspectives for acoustical tropical insect monitoring are promising. Considering “The Art of the Soluble” (Medawar 1967), we are convinced that progress in insect ecoacoustics will be accelerated by integrating traditional approaches with big data and citizen science initiatives.

## Supporting information

Soundscapes of figs 1,2

